# Differential effects of chronic hypertension and sex on vascular density along the septotemporal axis of the hippocampus in a rat model of VCID

**DOI:** 10.64898/2026.10.31.763336

**Authors:** Jared Luchetta, Emily D. Schellinger, Ziv Bolton, Gianna M. DeMarco, Abbie C. Chapman

**Author notes:** Correspondence to: Abbie C. Chapman, Ph.D. 4400 Vestal Parkway East Science 4 Building, Room 262 Binghamton, NY, USA 13902.

## Abstract

**Background:** Chronic hypertension is a major modifiable risk factor for vascular contributions to cognitive impairment and dementia (VCID), yet vascular changes that precede cognitive dysfunction remain poorly understood. This is particularly relevant for the hippocampus, a deep brain structure critical to memory function, which differs along its septotemporal axis functionally and in its susceptibility to hypoxic/ischemic injury. The impacts of hypertension and sex on hippocampal vascularization remains unclear and could contribute to the development/progression of VCID.

**Methods:** In this study, adult male and female Wistar and spontaneously hypertensive rats (SHR) were used at 5 months old, an age before memory impairments. Vascular density and morphometric parameters in dorsal and ventral hippocampus were quantified using high-resolution confocal microscopy and 3D vascular reconstruction.

**Results:** In SHR, vascular density was increased in the dorsal hippocampus in males and across dorsal and ventral hippocampus in females compared to normotensive rats. Diameter and length of the microvasculature shifted to be smaller and longer during hypertension in males, but not females.

**Conclusions:** These findings demonstrate chronic hypertension increases vascular density differentially by sex along the septotemporal axis of the hippocampus in mid adulthood that may increase vascular resistance and potentiate perfusion deficits or increase sites of blood-brain barrier disruption and contribute to VCID.

## Background

Vascular contributions to cognitive impairment and dementia (VCID) are the second major cause of dementia after Alzheimer’s disease and encompass a spectrum of cognitive dysfunction, with the most severe form being vascular dementia.^1–3^ Chronic hypertension is a particularly important contributor to VCID due to its high global incidence and association with dementia risk later in life. It affects approximately 1.4 billion people worldwide, making it the most prevalent of the 14 recognized modifiable dementia risk factors. More specifically, chronic hypertension is a primary cause of cerebrovascular disease and has the greatest impact on cognitive function when it develops early in adulthood.^4–8^ Chronic, uncontrolled hypertension in midlife increases the risk of dementia by ∼ 50% compared to normotensive individuals, highlighting the importance of understanding the impact of chronic hypertension on the cerebral circulation in order to combat VCID.^9–11^

Chronic hypertension contributes to cognitive decline by disrupting the cerebrovasculature and perfusion of brain regions particularly vulnerable to injury, such as the hippocampus.^12, 13^ In the healthy, state the hippocampus, a deep brain region central to learning and memory, has one of the lowest vascular densities in the brain and is substantially less vascularized than the cerebral cortex.^14, 15^ Hippocampal neurons are highly metabolically active and reliant upon blood flow to match such high demand.^16^ Despite lower vascularization, neuronal metabolic demand in the hippocampus remains sufficiently coupled with blood flow under healthy conditions.^14, 15, 17^ However, the limited vascular density leaves little reserve for reductions in perfusion and/or capillary rarefaction, which can occur during chronic hypertension. The current understanding of hypertension-induced cerebrovascular density changes comes largely from studies of the cerebral cortex in rodent models (e.g., spontaneously hypertensive rats, SHR) of the male sex. In male SHR, cortical vascular density findings are inconsistent in early adulthood (between 2 and 4 months of age), with studies reporting rarefaction, no change, or increased vascular density.^18^ As these studies have focused on the cortex, less is known about vascularization changes in the hippocampus. Similarly, these studies focused on males, and much less is known about females. Whether vascular density is uniform across the hippocampal septotemporal axis, subfields or between sexes – under healthy conditions or during hypertension – remains unclear.

The hippocampus is functionally diverse along its septotemporal axis. The dorsal and ventral hippocampus support different cognitive and emotional processes such that region-specific vascular density changes could carry distinct consequences. The dorsal hippocampus (posterior hippocampus in humans) is often studied due to its involvement in spatial memory, whereas the ventral hippocampus (anterior hippocampus in humans) is important for emotional processes (e.g., anxiety, stress) and is also disrupted during chronic hypertension.^19–22^ Although hypertension is more prevalent in men during midlife, women in general have a higher incidence of anxiety and stress-related disorders than men, functions closely tied to the ventral hippocampal circuitry.^23^ Together, these regional differences in hippocampal function suggest that the effect of hypertension may not be uniform across the dorsal and ventral hippocampus.

Given the critical role of the vasculature in tissue perfusion, nutrient/metabolite exchange, and vascular resistance, characterizing hypertension-induced changes in vascular density and morphometry in the hippocampus in mid-life may provide important information regarding its vulnerability and contributions to VCID.^3, 24^

In the current study, we evaluated hippocampal vascular architecture at micron-level resolution across the septotemporal axis in both sexes and whether hypertension impacted this vascular pattern that could contribute to VCID. Using lectin-based vascular labelling combined with high-resolution confocal microscopy, 3D hippocampal vascular reconstruction was used to determine vascularization of dorsal and ventral hippocampus. Vascular density and vessel sizes were assessed in hippocampal subregions (dentate gyrus, Cornu Ammonis, CA1, CA3) that could alter vascular resistance and hemodynamic parameters contributing to VCID.^25^ We hypothesized that vascular patterns would be similar between males and females across the septotemporal, and that chronic hypertension would cause capillary rarefaction throughout the dorsal and ventral hippocampus, including hippocampal subfields.

## Methods Animals

Male and female Wistar rats and SHR were purchased from Charles River at 3 months of age (N=20) and housed in the Binghamton University - State University of New York animal facility, an Association for Assessment and Accreditation of Laboratory Animal Care International (AAALAC) accredited facility. Rats were housed in groups with a tunnel/chew block and maintained on a 12-h light/dark cycle with ad libitum access to food and water. As this study did not include interventions/manipulations, rats were not randomly assigned to group.

SHR are a widely used model of chronic hypertension used to investigate hypertension-related cerebrovascular disease and share similar neuropathology of VCID.^26^ While Wistar-Kyoto rats are the genetic control for SHR, they are an established model of depression.^27–29^ Therefore, Wistar rats are often used in studies regarding age- and hypertension-related VCID and were used in the current study as normotensive controls.^30–32^ All procedures were approved by the Binghamton University Institutional Animal Care and Use Committee (IACUC Protocol #s 898-23 and 956-25; PI: Chapman) and conducted in accordance with the National Institutes of Health (NIH) Guide for the Care and Use of Laboratory Animals and according to the ARRIVE 2.0 guidelines. Euthanasia occurred via exsanguination under isoflurane anesthesia (3% in oxygen) followed by rapid decapitation.

### Vascular staining *in vivo*

At 5 months of age, rats were anesthetized with isoflurane (3% in oxygen), and tomato lectin (0.5 mL, 0.4 mg/mL in 0.1M PBS) was injected into the lateral tail vein and allowed to circulate for 5 minutes (n=5 rats/group). The circulation was flushed with ice-cold 0.1M PBS (pH 7.4) via transcardiac perfusion (60-80mL, 5mL/min). Brains were rapidly extracted from the skull and immersion fixed in cold 4% paraformaldehyde overnight (16-18 hours) at 4 °C and were then washed 3 times with 0.1 M PBS. Two coronal 300 μm thick serial sections of dorsal hippocampus were prepared with a Leica VT1200 vibratome. To match hippocampal morphology between regions, the brain was reoriented and secured to a wedge-shaped agarose block at a 75% incline and two 300 μm thick serial cross sections of ventral hippocampus taken. Brain sections were mounted with ProLong Diamond antifade (ThermoFisher, P36970) and then covered with a #1.5 superslip (Avantor, 48393-251). One female and two male Wistar rats were excluded from analyses due to technical issues resulting in incomplete tomato lectin perfusion. Multiple perfusions were completed on consecutive days to allow consistency of injections and tissue processing. Tomato lectin prepared daily and protected from light.

### High resolution confocal microscopy and quantification of vascular density

Images of microvasculature were acquired on a Leica DMi8 confocal microscope using hybrid detectors in standard mode. Z-stacks were captured through the entirety of the tissue slab at a z-step size of 3 microns with 1024x1024 pixel dimensions and a scan speed of 200 μm per second with a 10x Leica objective lens (Leica). Images were exported as .lif files and uploaded into Imaris developed by Oxford Instruments (v10.2). Whole hippocampus and hippocampal subfield regions of interest were generated using the Imaris Surfaces module. Subfield ROIs included the dentate gyrus, CA1 and CA3. For each reconstructed region, a standardized surface was first generated to define the region volume, after which the vasculature contained within that region was analyzed via an automated surfaces protocol. Vasculature was segmented using a seed diameter of 15.1 μm. Vessel segments were grouped into four volume bins: 0-10,000; 10,000-20,000, 0-20,000, and >20,000 μm^3^. For length analyses, vessels within the 0-20,000 μm^3^ range were isolated, and Imaris Filament tracer was used to quantify vessel length. In Filament analyses, length density was normalized to whole hippocampal volume from Imaris Surface analyses (length density = total vessel length / hippocampal volume). All images were analyzed using automation within Imaris and completed by two independent investigators that were blinded to group.

### Drugs and solutions

Tomato lectin DyLight 649 was purchased from Vector Laboratories (Newark, CA, USA). 0.1M PBS contained anhydrous sodium phosphate dibasic (Sigma Aldrich, S5136, 16.6 mM), anhydrous potassium phosphate monobasic (Sigma Aldrich, 71496, 3.9 mM), and NaCl (Fisher, BP358, 0.15 mM) and was made fresh weekly.

### Statistical analysis

A statistical power calculation was done to determine the number of animals needed using a two-sided 95% confidence interval for a single mean and 1 – β of 0.80 based on our studies quantifying microvascular density.^33^ Statistical analysis was performed using GraphPad Prism version 10.6.0 (GraphPad Software Inc., La Jolla, CA, USA). Comparisons were made to determine main effects and the interaction between two independent variables (sex and hypertension) using a two-way analysis of variance (ANOVA) with Tukey post-hoc tests to correct for multiple comparisons. Any data greater than two standard deviations from the mean were excluded from analyses, which was established *a priori*. To determine effects of brain region on vascular parameters within sex, a two-way mixed-effects model with a nested design and Sidak’s multiple comparison test was used. Sphericity was assumed, tested with Mauchly’s Test, and Greenhouse-Geisser correction used if needed. Data represent mean ± standard error of the mean (SEM) and significance was considered p-value < 0.05.

### Use of artificial intelligence (AI)

AI, including generative AI, were not used in any capacity for this manuscript, figures or tables.

## Results

To determine whether there were sex differences in vascularization of the hippocampus across the septotemporal axis, and how chronic hypertension may disrupt perfusion networks, we used systemic tomato lectin-DyLight649 labelling to visualize hippocampal vasculature in adult male and female Wistar rats and SHR. Figures 1A and 2A show representative images of the vascularization of cross sections of dorsal and ventral hippocampus from male and female Wistar rats and SHR. In the dorsal hippocampus, there was a main effect of hypertension on vascular density, with a trending effect of sex, and no interaction between the two (F_1,30_ = 16.05, p<0.001; F_1,30_ = 3.535, p=0.0698; F_1,30_ = 2.623, p=0.1158 by two-way ANOVA, respectively). Vascular density was increased during chronic hypertension in males (p<0.01 by two-way ANOVA with Tukey post-hoc test) but remained unchanged in females (Figure 1B). There were no differences in total vascular density between male and female Wistar rats (Figure 1B).

**Figure 1.**
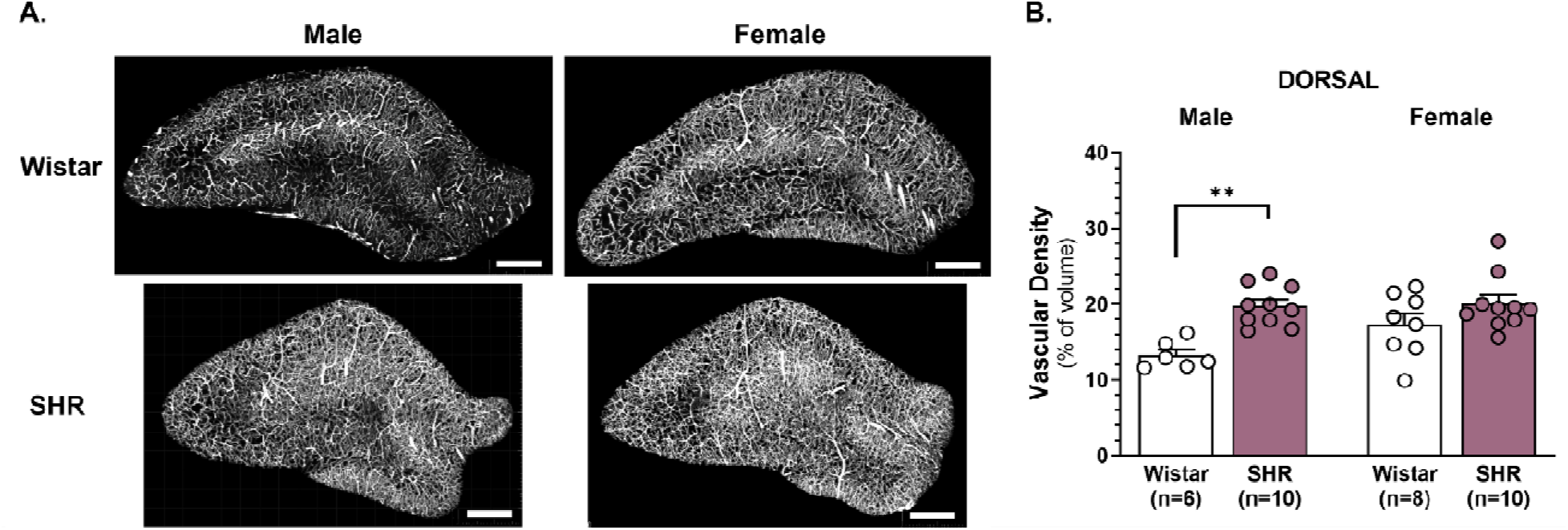
Vascular density in the dorsal hippocampus. **A.** Representative images of the vasculature in male and female dorsal hippocampus in both normotensive (Wistar) and hypertensive (SHR) animals **B.** Chronic hypertension increased vascular density in male, but not female, dorsal hippocampus. Each data point represents vascular density measured from one 300 μm thick hippocampal slab, with two slabs analyzed per animal. **p < 0.01. Data are presented as mean ± SEM and were analyzed by Two-Way ANOVA followed by Tukey’s post hoc test for multiple comparisons. n = 3 male Wistar, n= 5 male SHR, n = 4 female Wistar, and n = 5 female SHR. Scale bar = 500 μm.

**Figure 2.**
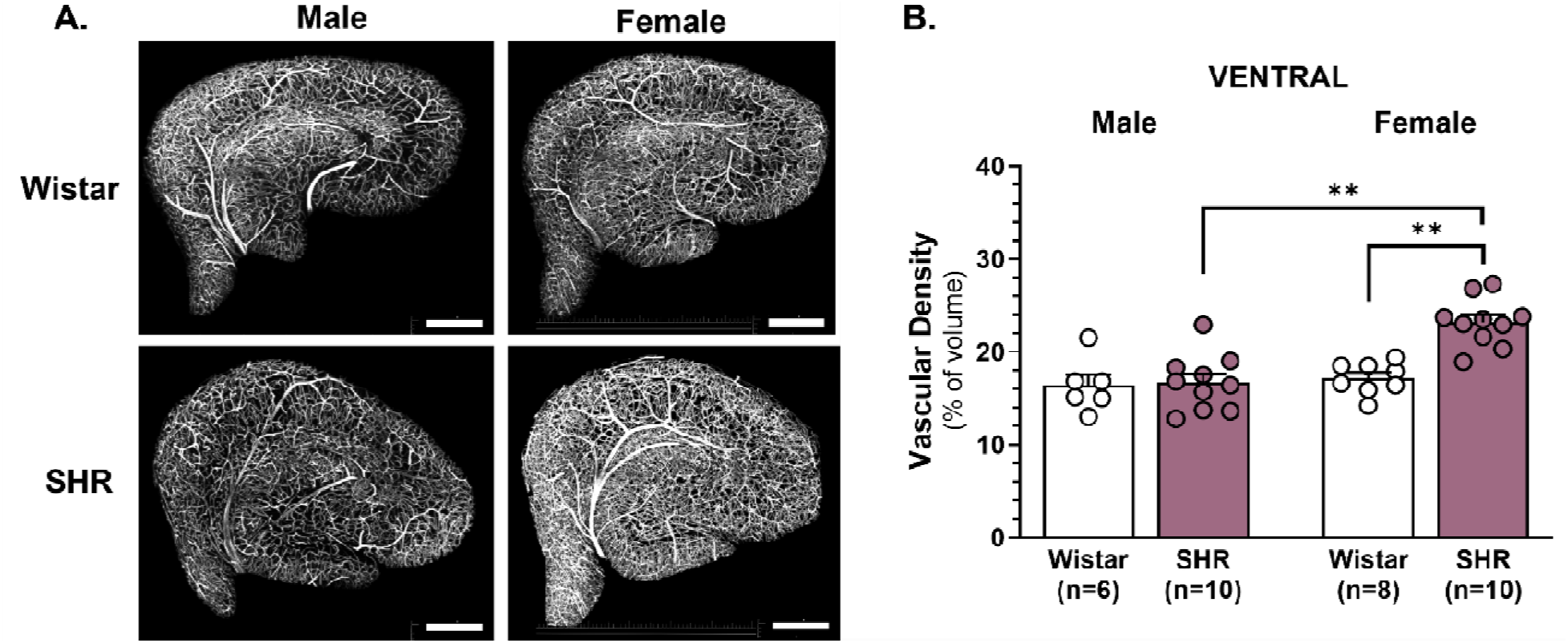
Vascular density in the ventral hippocampus. **A.** Representative images of the ventral hippocampus vasculature in male and female normotensive Wistar and hypertensive SHR. **B.** Chronic hypertension did not alter vascular density in male animals, but significantly increased vascular density female ventral hippocampus compared with female Wistar controls and male SHR. Each data point represents vascular density measured from one 300 μm thick hippocampal slab, with two slabs analyzed per animal. **p < 0.01. Data are presented as mean ± SEM and were analyzed by Two-Way ANOVA followed by Tukey’s post hoc test for multiple comparisons n = 3 male Wistar, n= 5 male SHR, n = 4 female Wistar, and n = 5 female SHR. Scale bar = 500 μm.

In ventral hippocampus, there were main effects of sex, hypertension, and a significant interaction between hypertension and sex on vascular density (F_1,30_ = 16.03, p<0.001; F_1,30_ = 11.91, p=0.0017; F_1,30_ = 9.732, p=0.0040 by two-way ANOVA, respectively). While there were no differences between normotensive males and females, ventral hippocampal vascular density was higher in female SHR compared to female Wistar and male SHR (p<0.01 by two-way ANOVA with Tukey post-hoc test; Figure 2B).

To understand potential regional differences in hippocampal vascularization, vascular density between dorsal and ventral hippocampus were compared in both male and female rats. In males, there was a main effect of chronic hypertension, no effect of region on hippocampal vascularization, and a significant interaction between hypertension and region (F_1,14_ = 9.545, p=0.0080; F_1,14_ = 0.00001, p=0.9969; F_1,14_ = 13.12, p=0.0028 by two-way mixed-effects analysis, respectively). In male SHR, the dorsal hippocampus had higher vascular density than ventral (p=0.0205 by two-way mixed-effects analysis with Šídák’s post-hoc test). In normotensive males, vascular density in dorsal hippocampus was lower in ventral hippocampus, but this did not reach statistical significance (p=0.0749 by two-way mixed-effects analysis with Šídák’s post-hoc test). In females, there was a main effect of hypertension, but no effect of region and no interaction between the two on vascularization (F_1,32_ = 16.78, p=0.0003; F_1,32_ = 1.948, p=0.1724; F_1,32_ = 2.324, p=0.1372 by two-way mixed-effects analysis, respectively). More specifically, there was no difference in vascular density between the dorsal and ventral hippocampus in female Wistar rats; however, the ventral hippocampus had higher vascularization compared to the dorsal hippocampus in female SHR, although this did not reach statistical significance (p=0.0705 by two-way mixed-effects analysis with Šídák’s post-hoc test).

Hippocampal subregions, including dentate gyrus, CA1, and CA3 have distinct functions and vary in susceptibility to hypoxic/ischemic injury that may be due, in part, to differences in vascular density.^14, 34^ To understand vascularization between hippocampal subregions, and whether this differs by sex and/or chronic hypertension, we compared vascular density between dentate gyrus, CA1 and CA3 of dorsal (Figure 3A) and ventral hippocampus (Figure 4A). In dorsal hippocampus, there were main effects of sex and hypertension, but no interaction between the two on vascular density of dentate gyrus (F_1,30_ = 6.487, p=0.0162; F_1,30_ = 16.57, p=0.0003; F_1,30_ = 0.2568, p=0.6161 by two-way ANOVA, respectively; Figure 3B) and CA1 (F_1,30_ = 9.633, p=0.0041; F_1,30_ = 23.47, p<0.0001; F_1,30_ = 0.4339, p=0.5151 by two-way ANOVA, respectively; Figure 3C). In CA3, there were main effects of both sex and hypertension and a significant interaction between sex and hypertension on vascular density (F_1,30_ = 6.106, p=0.0194; F_1,30_ = 23.85, p<0.0001; F_1,30_ = 4.218, p=0.0488 by two-way ANOVA, respectively; Figure 3D). There were no differences in vascularization of dentate gyrus and CA1 between males and females; however, chronic hypertension increased vascular density in both subregions (p<0.05 by two-way ANOVA with post-hoc Tukey test; Figures 3B and 3C). In CA3, male Wistar rats had lower vascular density compared to female Wistar rats and male SHR (p<0.05 by two-way ANOVA with post-hoc Tukey test; Figures 3D).

**Figure 3.**
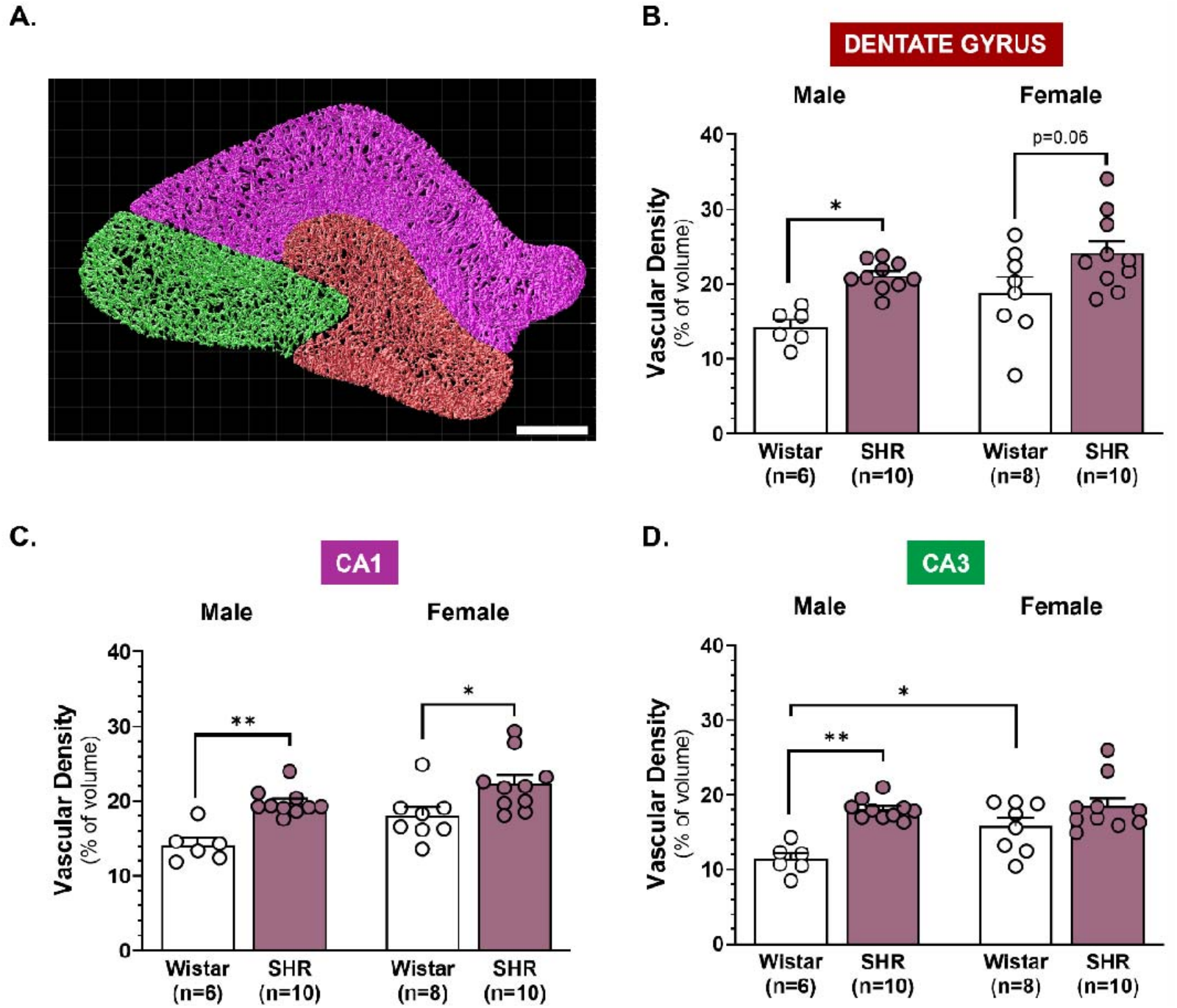
Vascular density in dorsal hippocampal subfields. **A.** Representative image identifying dentate gyrus (red), CA1 (magenta), and CA3 (green) regions of interest within the dorsal hippocampus. **B.** Chronic hypertension increased vascular density in male dentate gyrus, while females showed a nonsignificant trend toward increased vascular density. **C.** Chronic hypertension increased vascular density in CA1 of both males and females. **D.** Chronic hypertension increased vascular density in male CA3 compared to normotensive Wistar controls. Among Wistar animals, vascular density was greater in female than male CA3, whereas no difference observed between SHR and Wistar females. *p < 0.05, **p < 0.01. Data are presented as mean ± SEM and were analyzed by Two-Way ANOVA followed by Tukey’s post hoc test for multiple comparisons n = 3 male Wistar, n= 5 male SHR, n = 4 female Wistar, and n = 5 female SHR. Scale bar = 500 μm.

**Figure 4:**
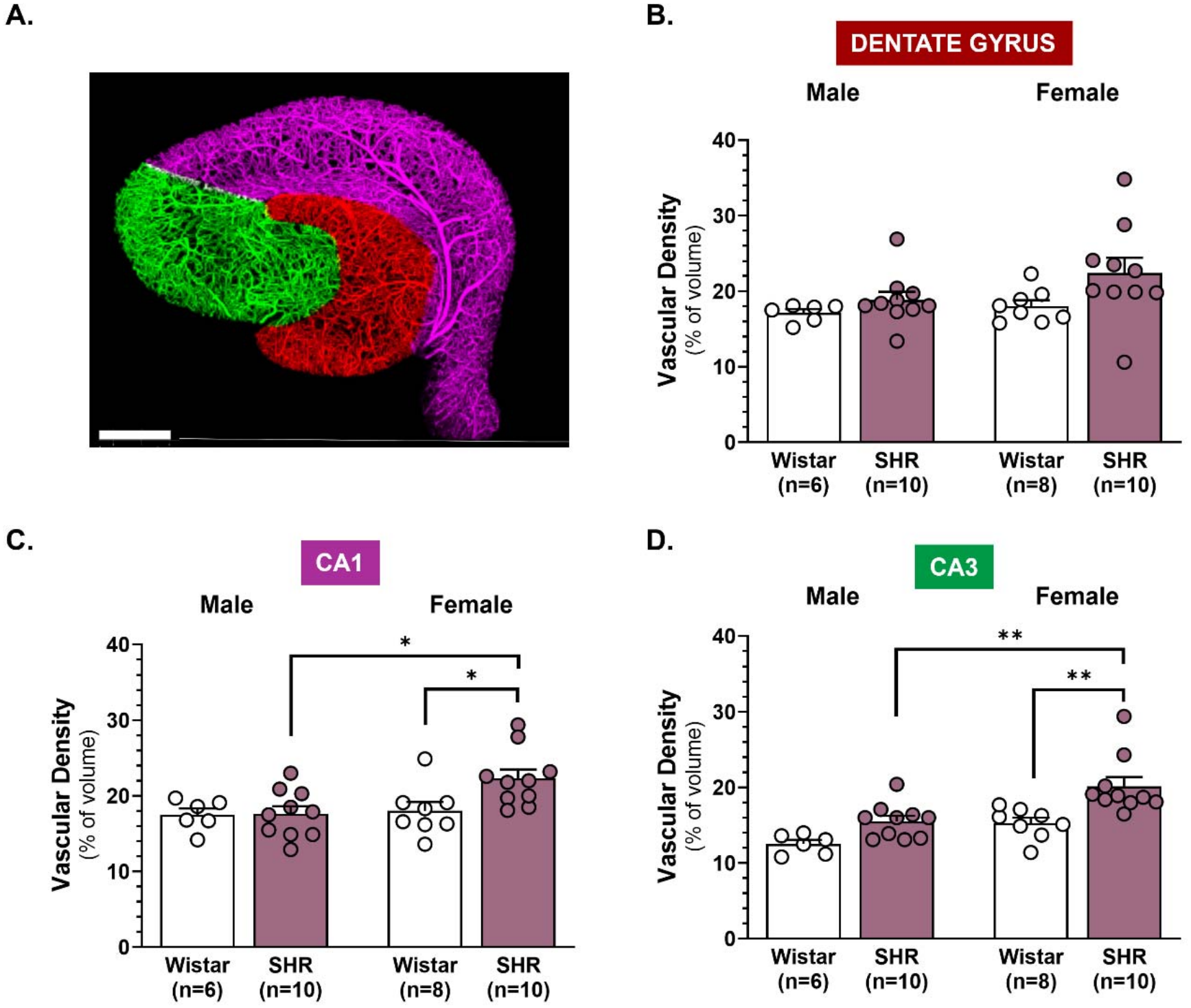
Vascular density in ventral hippocampal subfields. **A.** Representative image identifying Dentate Gyrus (red), CA1 (magenta), and CA3 (green) regions of interest within the <u>ventral</u> hippocampus. **B.** Vascular density was unchanged in the dentate gyrus of male and females **C.** Chronic hypertension did not change vascular density in male CA1 vascular density in males but increased vascular density in females compared to female Wistar and male SHR. **D.** Chronic hypertension did not alter CA3 vascular density in males, whereas vascular density was increased in female SHR compared to female Wistar and male SHR. *p < 0.05, **p < 0.01. Data are presented as mean ± SEM and were analyzed by Two-Way ANOVA followed by Tukey’s post hoc test for multiple comparisons. n = 3 male Wistar, n= 5 male SHR, n = 4 female Wistar, and n = 5 female SHR. Scale bar = 500 μm.

In ventral hippocampus, there was a main effect of hypertension, but no effect of sex or interaction between the two on vascular density in the dentate gyrus (F_1,30_ = 4.469, p=0.0429; F_1,30_ = 2.386, p=0.1329; F_1,30_ = 0.8463, p=0.3649 by two-way ANOVA, respectively; Figure 4B); however, no group differences were detected with Tukey’s post-hoc test (p>0.05). In CA1, there was a main effect of sex, but not hypertension on vascular density and no interaction between the two (F_1,30_ = 5.213, p=0.0297; F_1,30_ = 3.786, p=0.0611; F_1,30_ = 3.407, p=0.0748 by two-way ANOVA, respectively; Figure 4C). In the CA3, there were main effects of sex and hypertension on vascularization but no interaction between sex and hypertension (F_1,30_ = 15.35, p=0.0005; F_1,30_ = 17.43, p=0.0002; F_1,30_ = 0.9903, p=0.3276 by two-way ANOVA, respectively; Figure 4D). Female SHR had higher vascular density compared to female Wistar and male SHR in both CA1 and CA3 subregions of ventral hippocampus (p<0.05 by two-way ANOVA with post-hoc Tukey test; Figures 4C and 4D). Together, these data suggest that there were no baseline sex differences in vascularization of dorsal or ventral hippocampus, chronic hypertension differentially affected vascular density in a sex- and region-dependent manner.

Greater vascular density could reflect a shift toward larger vascular structures, an increased prevalence of small vascular structures, or a combination of both, with each pattern carrying distinct implications for local hemodynamics. Given the region- and sex-dependent differences in overall vascular density, we next determined whether these differences were associated with changes in vessel size in dorsal or ventral hippocampus. Figure 5A shows representative images of the cross-section of dorsal hippocampus (left) and the vascular segment volume heatmap (right) used to quantify vessel size, with cooler colors being smaller and warmer colors being larger. In the dorsal hippocampus there was a main effect of hypertension, but not sex, and a significant hypertension-by-sex interaction on mean segment volume (F_1,28_ = 13.16, p=0.0011; F_1,28_ = 1.203, p=0.2822; F_1,28_ = 8.443, p=0.0071 by two-way ANOVA, respectively; Figure 5B). In ventral hippocampus, there was a main effect of hypertension, but no effect of sex on mean segment volume, and no interaction between the two (F_1,30_ = 8.885, p=0.0057; F_1,30_ = 1.414, p=0.2438; F_1,30_ = 1.366, p=0.2517 by two-way ANOVA, respectively; Figure 5C). There were no sex differences between mean segment volume in dorsal or ventral hippocampus; however, segment volume was significantly lower in male SHR compared to normotensive male rats in both regions, with hypertension having no effect on vessel size in females (p<0.01 by two-way ANOVA with Tukey’s post-hoc test).

**Figure 5:**
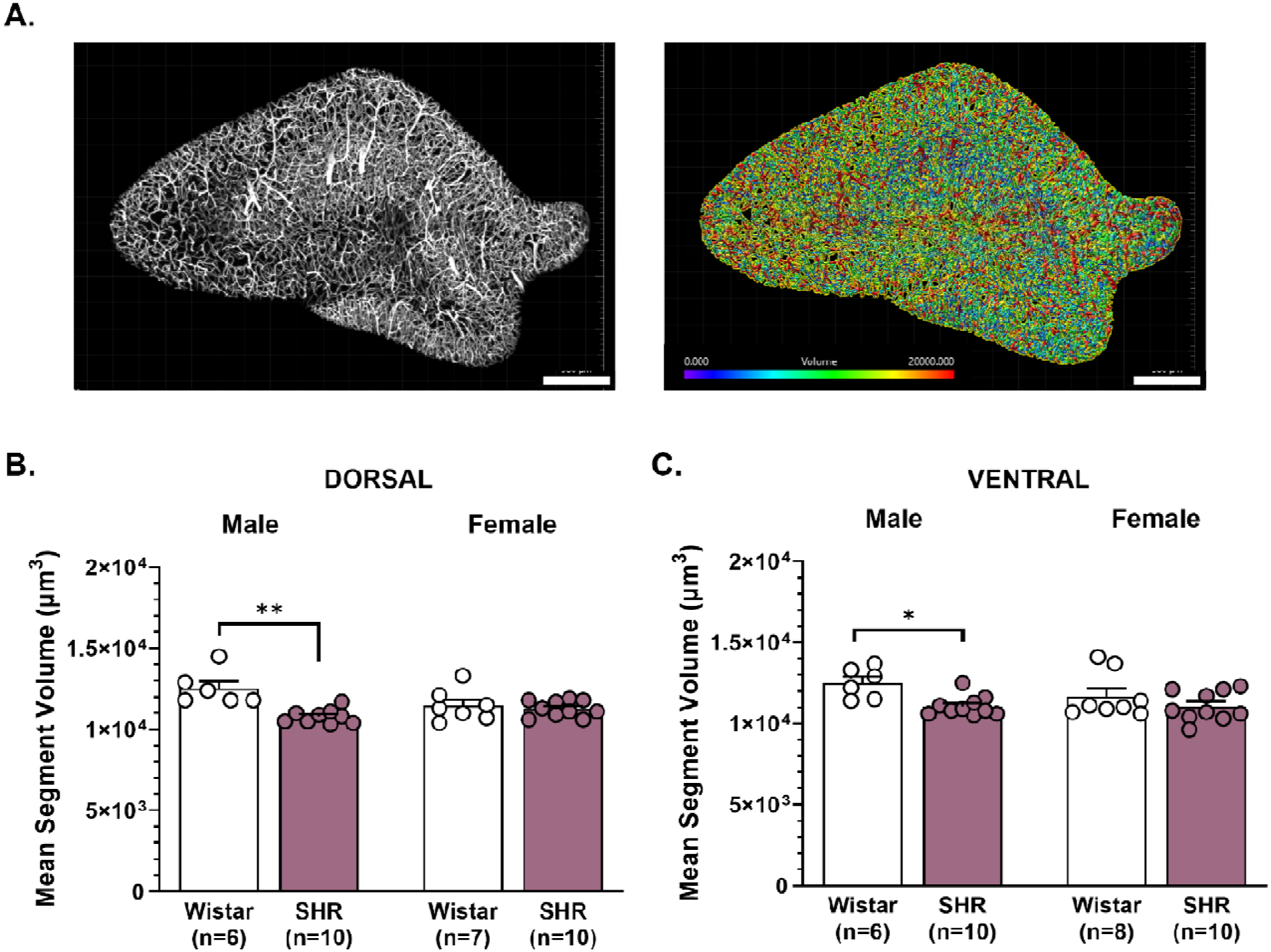
Hypertension decreases mean vessel segment volume in males. **A.** Representative image of hippocampal vasculature (left) and 3D-reconstructed, segmented vasculature (right), with vessel segments color coded by size; red indicates larger vessel segments and blue indicates small vessel segments. **B.** Chronic hypertension decreased mean segment volume in male dorsal hippocampus, whereas no change was observed in females. **C.** Chronic hypertension also decreased mean segment volume in male ventral hippocampus, with no change observed in females. *p < 0.05. Data are presented as mean ± SEM and were analyzed by Two-Way ANOVA followed by Tukey’s post hoc test for multiple comparisons. n = 3 male Wistar, n= 5 male SHR, n = 4 female Wistar, and n = 5 female SHR. Scale bar = 500 μm.

To understand if there were regional differences between overall vessel size and if chronic hypertension impacted this, we compared mean segment volume between dorsal and ventral hippocampus in both sexes. In males, chronic hypertension was associated with smaller vasculature in general, as there was a main effect of hypertension on vessel size, with no effect of region or a significant interaction between region and hypertension (F_1,27_ = 33.01, p<0.0001; F_1,27_ = 0.1848, p=0.6707; F_1,27_ = 0.3035, p=0.5862 by two-way mixed effects analysis, respectively). In females, there were no main effects of hypertension or region on mean segment volume, and no interaction between the two (F_1,16_ = 1.376, p = 0.2575; F_1,15_ = 0.0989, p=0.7575; F_1,15_ = 0.3620, p=0.5564 by two-way mixed effects analysis, respectively). When stratified by hippocampal subregion, mean segment volume was consistently lower in male SHR compared to male Wistar in all subregions of dorsal hippocampus and in dentate gyrus and CA3 of ventral hippocampus (Table 1). There were no differences in mean segment volume in either dorsal or ventral hippocampal subregions between normotensive and hypertensive females (Table 1). One female Wistar rat was excluded from mean segment volume of dorsal hippocampus due to being a statistical outlier (> 2SD from the mean). These data suggest that chronic hypertension-induced reductions in mean segment volume occurred exclusively in males and indicate a shift toward smaller vascular structures in both dorsal and ventral hippocampus.

**Table 1:**
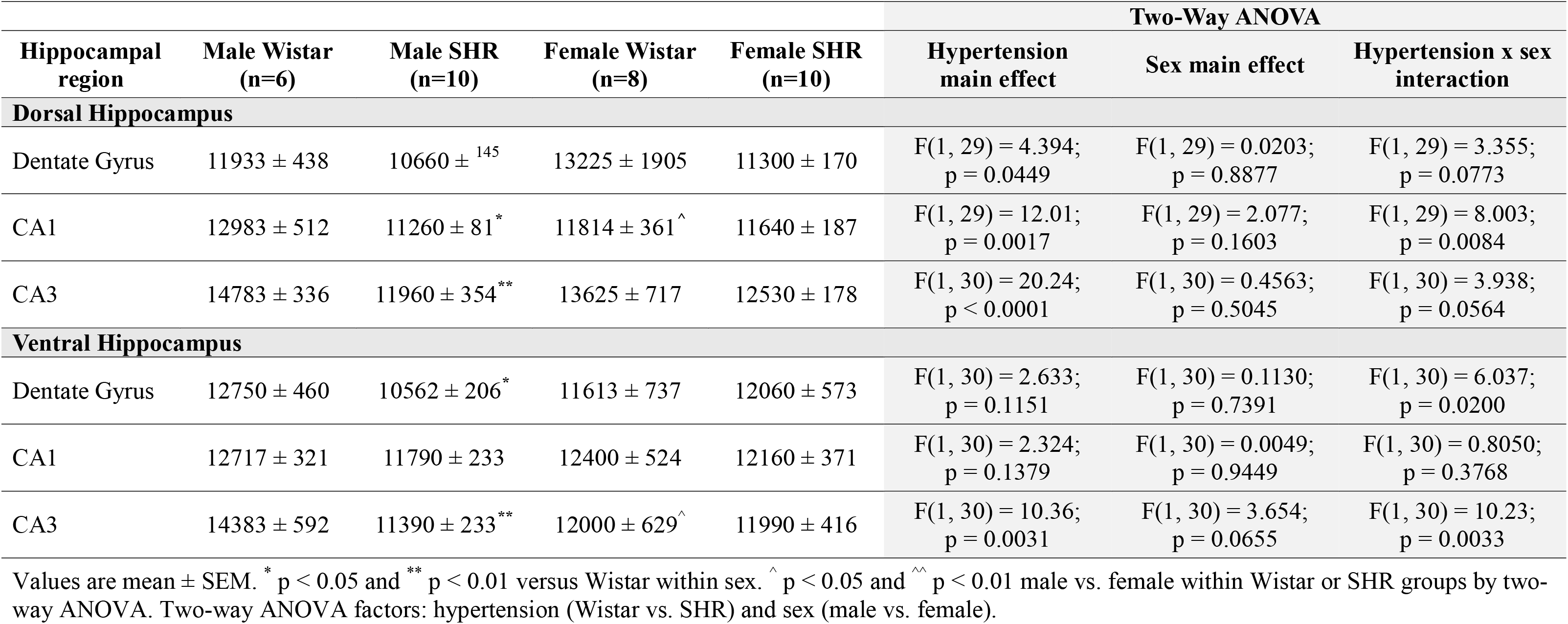
Mean vascular segment size in dorsal and ventral hippocampal subregions of male and female Wistar rats and SHR.

| Hippocampal region | Male Wistar (n=6) | Male SHR (n=10) | Female Wistar (n=8) | Female SHR (n=10) | Two-Way ANOVA |  |  |
|---|---|---|---|---|---|---|---|
|  |  |  |  |  | Hypertension main effect | Sex main effect | Hypertension x sex interaction |
| **Dorsal Hippocampus** |  |  |  |  |  |  |  |
| Dentate Gyrus | 11933 ± 438 | 10660 ± <sup>145</sup> | 13225 ± 1905 | 11300 ± 170 | F(1, 29) = 4.394;<br>p = 0.0449 | F(1, 29) = 0.0203;<br>p = 0.8877 | F(1, 29) = 3.355;<br>p = 0.0773 |
| CA1 | 12983 ± 512 | 11260 ± 81<sup>*</sup> | 11814 ± 361<sup>^</sup> | 11640 ± 187 | F(1, 29) = 12.01;<br>p = 0.0017 | F(1, 29) = 2.077;<br>p = 0.1603 | F(1, 29) = 8.003;<br>p = 0.0084 |
| CA3 | 14783 ± 336 | 11960 ± 354<sup>**</sup> | 13625 ± 717 | 12530 ± 178 | F(1, 30) = 20.24;<br>p < 0.0001 | F(1, 30) = 0.4563;<br>p = 0.5045 | F(1, 30) = 3.938;<br>p = 0.0564 |
| **Ventral Hippocampus** |  |  |  |  |  |  |  |
| Dentate Gyrus | 12750 ± 460 | 10562 ± 206<sup>*</sup> | 11613 ± 737 | 12060 ± 573 | F(1, 30) = 2.633;<br>p = 0.1151 | F(1, 30) = 0.1130;<br>p = 0.7391 | F(1, 30) = 6.037;<br>p = 0.0200 |
| CA1 | 12717 ± 321 | 11790 ± 233 | 12400 ± 524 | 12160 ± 371 | F(1, 30) = 2.324;<br>p = 0.1379 | F(1, 30) = 0.0049;<br>p = 0.9449 | F(1, 30) = 0.8050;<br>p = 0.3768 |
| CA3 | 14383 ± 592 | 11390 ± 233<sup>**</sup> | 12000 ± 629<sup>^</sup> | 11990 ± 416 | F(1, 30) = 10.36;<br>p = 0.0031 | F(1, 30) = 3.654;<br>p = 0.0655 | F(1, 30) = 10.23;<br>p = 0.0033 |

Values are mean ± SEM. <sup>\*</sup> p < 0.05 and <sup>\*\*</sup> p < 0.01 versus Wistar within sex. <sup>^</sup> p < 0.05 and <sup>^^</sup> p < 0.01 male vs. female within Wistar or SHR groups by two-way ANOVA. Two-way ANOVA factors: hypertension (Wistar vs. SHR) and sex (male vs. female).

To determine whether hypertension preferentially affected a specific segment of the vascular network, and if this differed by sex or hippocampal region, we divided the total vasculature into three size ranges based upon volume: the smallest vascular segments ≤ 10,000 µm^3^, vessels 10,000 – 20,000 µm^3^ and larger vascular segments > 20,000 μm^3^ (Figure 6A). As the dorsal hippocampus was affected in males and the ventral hippocampus in females, volumetric analyses focused only on these regions. In males, there were main effects of hypertension and segment size on vascular density in dorsal hippocampus, and a significant interaction between the two (F_1,14_ = 26.12, p = 0.0002; F_1.36, 19.05_ = 26.56, p<0.0001; F_1.36, 19.05_ = 11.84, p=0.0013 by two-way mixed effects analysis, respectively; Figure 6B). SHR had a higher density of all sizes of vessels compared to Wistar rats (p<0.01 by two-way mixed effects analysis with Šídák’s post-hoc test). In Wistar rats, there was a lower density of vasculature of the smallest volume compared to a similar density of the larger segments. However, in SHR, the density of vasculature of the mid-sized vessels was higher than both the smallest and largest vascular segments studied (p<0.01 by two-way mixed effects analysis with Šídák’s post-hoc test; Figure 6B). In females, there were main effects of hypertension and segment size on vascular density in the ventral hippocampus, but no interaction between the two (F_1,16_ = 33.38, p <0.0001; F_1.14, 18.2_ = 14.54, p=0.0009; F_1.14, 18.2_ = 0.9062, p=0.3668 by two-way mixed effects analysis, respectively; Figure 6C). SHR had a higher density of all sizes of vessels compared to Wistar rats (p<0.05 by two-way mixed effects analysis with Šídák’s post-hoc test). In Wistar rats, there was a lower density of vasculature of the smallest volume compared to vasculature of 10,000 – 20,000 µm^3^ (p<0.01 by two-way mixed effects analysis with Šídák’s post-hoc test) and no difference between density of the largest vascular segments. In female SHR, the distribution of vascular volumes was of a similar pattern as what occurred in males, and density of vasculature of the mid-sized vessels was higher than both the smallest and largest vascular segments studied (p<0.01 by two-way mixed effects analysis with Šídák’s post-hoc test; Figure 6C).

**Figure 6:**
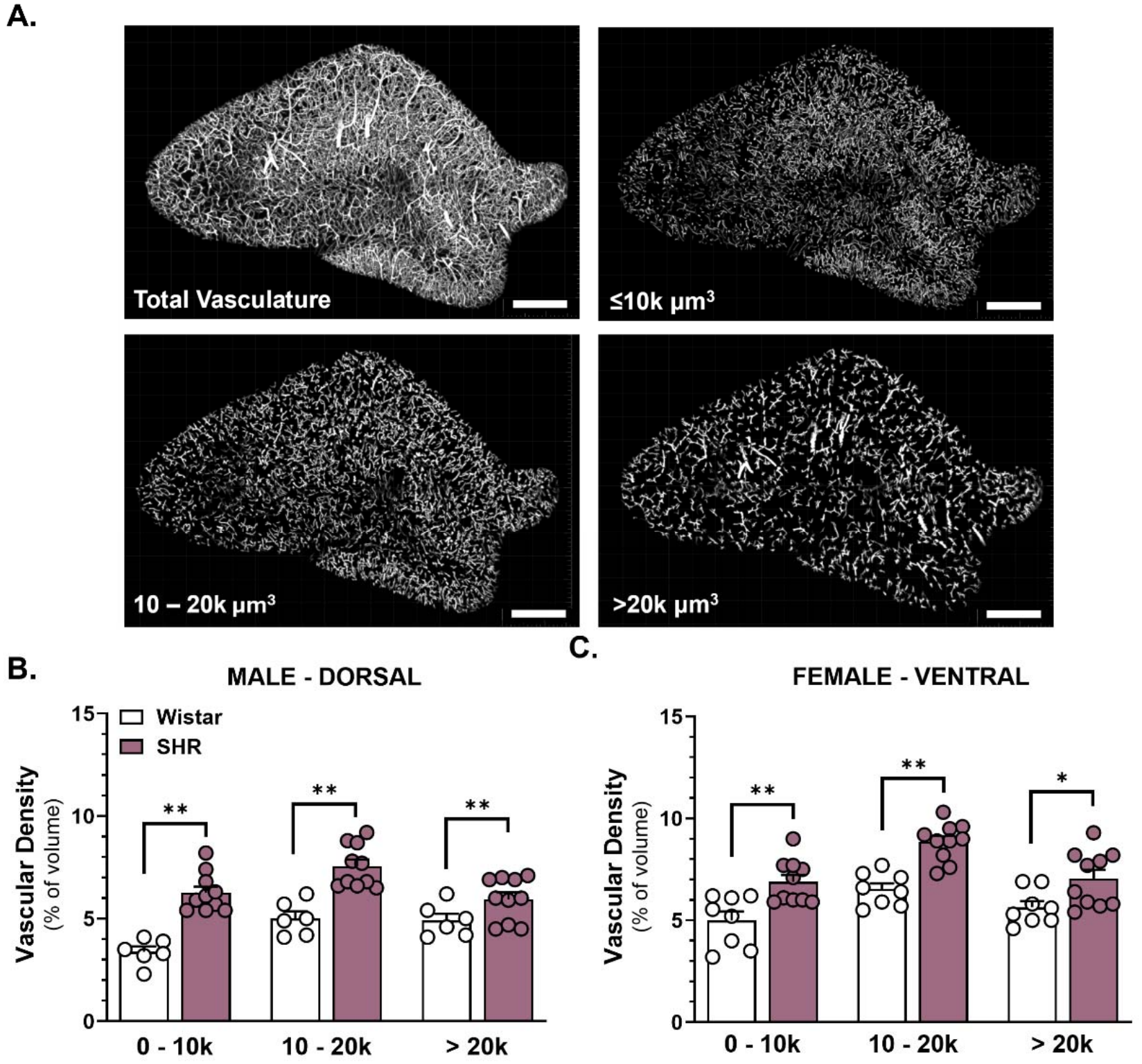
Hypertension increases vascular density across all vessel sizes. **A.** Representative masks showing total hippocampal vasculature (top left) and vascular segments stratified by volume (≤10,000 μm^3^, 10,000-20,000 μm^3^, and > 20,000 μm^3^). **B.** In the male dorsal hippocampus, vascular density was significantly increased in SHR compared with Wistar rats across all vessel size categories. **C.** In the female ventral hippocampus, vascular density was increased across all vessel size categories. *p < 0.05, **p < 0.01. Data are presented as mean ± SEM and were analyzed by Two-Way ANOVA followed by Tukey’s post hoc test for multiple comparisons. n = 3 male Wistar, n= 5 male SHR, n = 4 female Wistar, and n = 5 female SHR. Scale bar = 500 μm.

To determine whether the hypertension-induced shifts within the hippocampal microvasculature (≤ 20,000 μm^3^) were accompanied by changes in vascular morphology that contribute to vascular resistance and perfusion, we quantified mean microvessel length and length density in dorsal and ventral hippocampus. In the dorsal hippocampus, male SHR had increased mean length and higher length density compared to normotensive male rats, with there being no changes in morphological parameters during hypertension in the female sex (Table 2). In the ventral hippocampus, chronic hypertension did not affect mean vessel diameter or length density; however, mean length of microvasculature was larger in male SHR compared to male Wistar rats and female SHR (Table 2). There were no differences between sex in normotensive rats in either parameter studied.

**Table 2:**
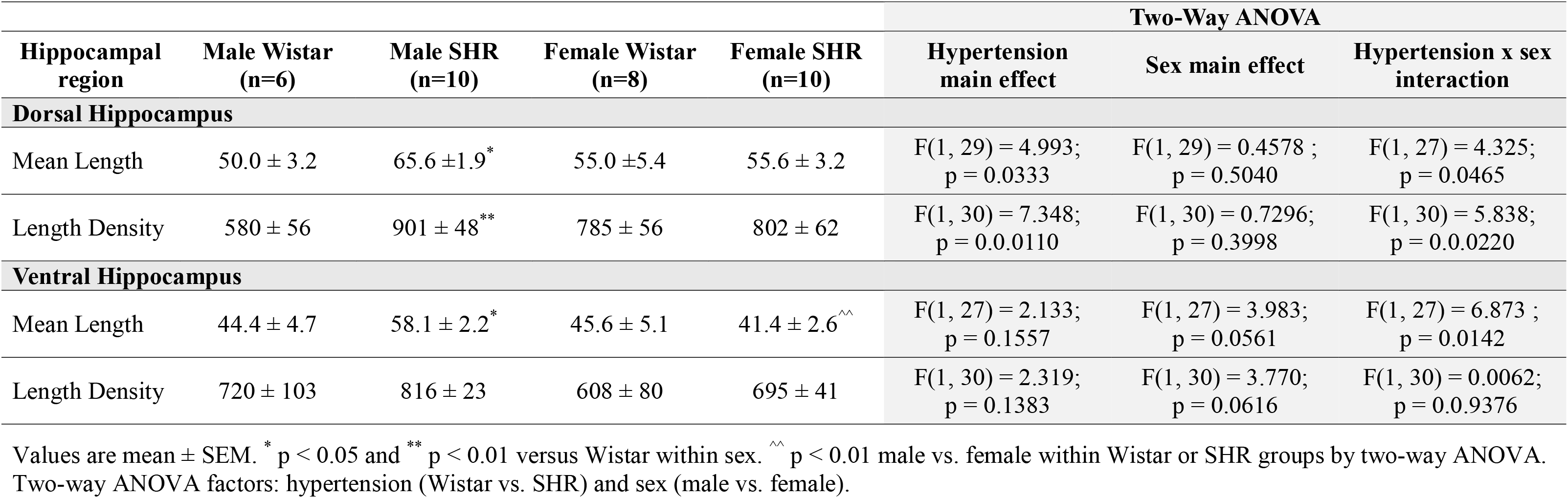
Vascular segment length and length density in the dorsal and ventral hippocampus.

## Discussion

Chronic hypertension is the leading cause of cerebrovascular disease affecting nearly 1.4 billion adults worldwide and is the most prevalent modifiable risk factor for VCID.^9^ Brain regions involved in memory function such as the hippocampus are directly damaged during chronic hypertension, resulting in arteriolar dysfunction, decreased blood flow and memory impairment that progress with age.^30^ How the structure of the vascular network changes in the hippocampus in early adulthood, when uncontrolled hypertension initiates progressive vascular injury, and whether one sex or hippocampal region/subregion is more affected remains unclear, and could contribute to VCID. Here, we found that chronic hypertension restructured the hippocampal vascular network differentially, depending upon sex and hippocampal region.

Contrary to our hypothesis that hypertension would be associated with vascular rarefaction in the dorsal and ventral hippocampi, chronic hypertension predominantly increased vascular density in the dorsal hippocampus of males and ventral hippocampus of females. Further, there was a shift towards smaller and longer microvessels specifically in males, although density was increased in all vascular sizes studied. Subfield-level analysis revealed that vascular adaptations occurred within specific hippocampal subfields even when overall regional vascular density was unchanged. These findings demonstrate that the hippocampal vascular network undergoes sex-, region- and subfield-specific increases in vascular density that may profoundly change vascular resistance and perfusion of this ischemia-prone brain region critical to memory.

Hippocampal blood flow and vascular reserve are recognized as critical factors in maintaining neurocognitive health later in life.^3, 30, 35^ Vascular resistance increases proportionately with vessel length (Resistance ∝ L, where L is vessel length) and rises steeply as vessel diameter decreases (Resistance ∝ 1/r^4^, where r is the vessel radius).^36^ Upstream of the capillary bed, penetrating arterioles and pial arteries are primary regulators of cerebrovascular resistance, and relatively small changes in diameter can substantially influence downstream perfusion. Alterations within the architecture of the microvascular network can further modify the distribution and organization of flow along the vascular tree and perfuse the hippocampal formation.^24, 37^ In the current study, the dorsal hippocampus in male and the ventral hippocampus female SHR had greater vascular density across all vessel sizes investigated, suggesting that there is a general increase in vascularization in early adulthood rather than expansion of specific segments along the vascular tree (i.e., capillary density). In males, decreased mean segment volume occurred alongside increased vascular density, suggesting a shift toward a greater proportion of microvessels. Further, this coincided with increased mean segment length and length density, although tortuosity was not assessed. The combination of increased vascular density, increased length density and reduced mean segment volume suggests there is increased vascular resistance in the dorsal hippocampus of males during chronic hypertension. However, we have previously shown that absolute blood flow in dorsal hippocampus was similar between normotensive and hypertensive male rats in early adulthood.^30^ Together, these data suggest that increased vascularization may occur as an adaptation to chronic hypertension to maintain appropriate blood flow. A similar paradoxical increase in cerebrovascular density has recently been reported in a transgenic mouse model of Alzheimer’s disease using longitudinal two-photon imaging.^38^ In early-to-mid adulthood, cerebrovascular density was substantially increased that corresponded with upregulation of genes involved in pro-angiogenic responses to hypoxia, suggesting a vascular adaptation to hypoperfusion. That increase in cerebral vascularization was transient and followed by decreased vascular density that progressed as a function of age.^38^ It remains unclear whether the hypertension-induced changes to the vascular network of the hippocampus found in the current study are present from birth or occur in response to hypoxia.

Hippocampal vascular architecture is not homogenous across the hippocampal formation, and its regional and subfield organization differs substantially from the vast microvascular networks in the neocortex (e.g., parietal cortex). Compared to the neocortex, the hippocampus has lower capillary density, lower resting blood flow and lower oxygenation, creating a relatively constrained vascular environment for meeting metabolic demand.^15, 39–45^ In addition to differences in the vascular architecture in the hippocampus, perfusion also differs by subfields. A recent study using high resolution 7T imaging reported that blood flow in CA1 was lower than CA3 and dentate gyrus.^46^ Importantly, CA1 is the most susceptible subfield to hypoxic/ischemic injury.^34^ Therefore, even relatively small changes in perfusion could have a tremendous impact on neuronal health and memory function. In the current study, we show increased vascular density in dorsal CA1 during chronic hypertension in both sexes. While increased vascular density could be protective, specifically at the capillary level, if the substantial increase in total vascularization reported in the current study persists, it could result in a profound increase in vascular resistance, potentiate perfusion deficits and cause neuronal injury over time. In addition, chronic hypertension is known to damage the blood-brain barrier (BBB), a highly specialized interface regulating the passage of systemically circulating factors into the brain.^32, 47^ The hypertension-induced increased hippocampal vascular density would provide greater surface area and potential for BBB disruption that could further contribute to neuroinflammation, neuronal injury and VCID.^48^

Along the septotemporal axis, the dorsal hippocampus is preferentially involved in spatial learning and memory, whereas the ventral hippocampus is associated with anxiety, fear, and stress-related behavior.^19, 22, 49^ Within these regions, individual subfields also support distinct computations, such as contextual discrimination, memory processing, pattern representation, spatial representation, fear/anxiety discrimination and approach-avoidance behavior.^49–52^ In our study, males exhibited increased vascular density throughout the dorsal hippocampus, including DG, CA3, and CA1, whereas females demonstrated a more selective pattern, with increased dorsal CA1 and ventral CA1 and CA3. The predominantly dorsal pattern of vascularization in males suggests metabolic support of neurocircuitry involved in spatial learning and memory, whereas the mixed dorsal/ventral pattern in females spans both spatial and stress/anxiety-related domains. Thus, hypertension does not produce a uniform change in vascularization across the septotemporal axis of the hippocampus, but instead preferentially alters vascular networks within specific functional domains. If the hypertension-related increased vascularization is indeed protective of perfusion and neuronal health early in adulthood, then susceptibility to hypoxic/ischemic injury and memory domain dysfunction during chronic hypertension depends upon sex. Females may be more protected in general, as vascularization was higher in subregions in both dorsal and ventral hippocampus, whereas males may undergo greater disruption in ventral hippocampus-dependent functions (e.g., anxiety). Time-course studies are needed to determine the relationships between hippocampal perfusion, neuronal health, cognitive function and vascular density across the lifetime during hypertension to better understand its potential role in progression of VCID in both sexes.

The distinct region-dependent vascularization patterns that emerge between males and females during chronic hypertension suggest that biological mechanisms regulating angiogenesis and rarefaction differ by sex. Differences in hypertensive burden, vascular protection, regional cerebral blood flow demand or vascular development could contribute to divergent responses between males and females. ^53–56^ Transcriptomic studies demonstrate sex-dependent differences in gene expression among cells in the neurovascular unit, distinct age-related changes in gene expression across the lifespan, and differential expression along the dorsal and ventral hippocampal axis.^57–59^ Gonadal hormones may provide an additional source of divergence, as estrogens, progesterone, and androgens can influence endothelial function, vascular tone, inflammatory signaling, and angiogenesis.^54, 56, 60^ The estrous cycle is known to influence functional aspects of the cerebrovasculature (e.g., endothelial-dependent vasodilation, cerebral blood flow) but not at the structural level, likely due to its rapid cycle, lasting only 4 days with some stages only lasting a few hours.^61, 62^ However, an ovariectomy reduces cerebral vascular density, an effect that is rescued by estrogen replacement.^63^ As VCID is a disease of aging, future studies are needed to determine how menopause affects vascularization of the hippocampus when cerebrovascular protection by estrogens is lost. Together, sex-dependent differences in neurovascular biology, including transcriptional regulation, vascular physiology, and hormone signaling, may contribute to regional patterns of hippocampal vascular adaptation during hypertension.

There are several limitations to the current study. Vascular morphometric analyses were completed during chronic hypertension in mid-adulthood. This is an age prior to onset of hypertension-induced hippocampal hypoperfusion and memory decline, at least as determined to occur in males.^30^ However, we were unable to determine whether increases in vascular density represent an early adaptive response, a stable phenotype or a transient stage that precedes rarefaction. Further, our measurements were structural and therefore cannot establish whether increased vascular density may be accompanied by changes in blood flow, vascular inflammation, neutrophil stalling or oxygen/glucose delivery.

## Conclusions

Chronic hypertension increased hippocampal vascular density in a sex-and region-specific manner, revealing distinct vascular responses in males and females. These findings suggest that vulnerability to VCID may be shaped by how the hippocampal vascular network changes across regions, between sexes and over time. This study advances our understanding of sex-specific changes in hippocampal vascular architecture during chronic hypertension that may contribute to sex-specific cognitive dysfunction in VCID.

## Abbreviations

ANOVA: analysis of variance
AI: artificial intelligence
AAALAC: Association for Assessment and Accreditation of Laboratory Animal Care International
BBB: blood-brain barrier
CA1: Cornu Ammonis 1
CA3: Cornu Ammonis 3
IACUC: Institutional Animal Care and Use Committee NIH: National Institutes of Health
PBS: phosphate buffered saline
SEM: standard error of the mean
SHR: spontaneously hypertensive rat
VCID: vascular contributions to cognitive impairment and dementia

## Declarations

### Ethics approval and consent to participate

All procedures were approved by the Binghamton University - SUNY Institutional Animal Care and Use Committee (IACUC Protocol #s 898-23 and 956-25; PI: Chapman) and conducted in accordance with the NIH Guide for the Care and Use of Laboratory Animals and according to the ARRIVE 2.0 guidelines

### Consent for publication

Not applicable.

### Availability of data and materials

All datasets generated and/or analyzed during the current study are publicly available within this manuscript and in the Harvard Dataverse repository (https://doi.org/10.7910/DVN/8OYELB). Due to the nature and size of the confocal images, these will be made available upon reasonable request.

### Authors’ contributions

JL was involved in the study design, *in vivo* tomato lectin infusions, tissue preparation, imaging, image analyses, statistical analyses, constructing figures, writing the initial draft of the manuscript. EDS, ZB and GMD were involved in tissue preparation, imaging and image analyses. ACC was involved in conceptualization, study design, funding acquisition, supervision, resources, statistical analyses, constructing figures, and writing the initial draft of the manuscript. All Authors read and approved the final manuscript.

## Acknowledgements

Not applicable.

